# Aberrant connectivity in auditory precision encoding in schizophrenia spectrum disorder and across the continuum of psychotic-like experiences

**DOI:** 10.1101/792473

**Authors:** Kit Melissa Larsen, Ilvana Dzafic, Hayley Darke, Holly Pertile, Olivia Carter, Suresh Sundram, Marta I. Garrido

## Abstract

**Background:** The ability to generate a precise internal model of statistical regularities is impaired in schizophrenia. Predictive coding accounts of schizophrenia suggest that psychotic symptoms may be explained by a failure to build precise beliefs or a model of the world. The precision of this model may vary with context. For example, in a noisy environment the model will be more imprecise compared to a model built in an environment with lower noise. However compelling, this idea has not yet been empirically studied in schizophrenia. Methods: In this study, 62 participants engaged in a stochastic mismatch negativity paradigm with high and low precision. We included inpatients with a schizophrenia spectrum disorder (N=20), inpatients with a psychiatric disorder but without psychosis (N=20), and healthy controls (N=22), with comparable sex ratio and age distribution. Bayesian mapping and dynamic causal modelling were employed to investigate the underlying microcircuitry of precision encoding of auditory stimuli. Results: We found strong evidence (exceedance p > 0.99) for differences in the underlying connectivity associated with precision encoding between the three groups as well as on the continuum of psychotic-like experiences assessed across all participants. Critically, we show changes in interhemispheric connectivity between the two inpatient groups, with some connections further aligning on the continuum of psychotic-like experiences. Conclusions: While our results suggest continuity in backward connectivity alterations with psychotic-like experiences regardless of diagnosis, they also point to specificity for the schizophrenia spectrum disorder group in interhemispheric connectivity alterations.

## Introduction

The ability to adapt to the ever changing environment and make inferences about future events is robustly reduced in people with schizophrenia^1–3^. This has been studied in a classical laboratory setting using auditory oddball paradigms. In auditory oddball paradigms, unpredictable (“oddball”) sounds are interspersed in a stream of frequently occurring standard sounds. Such paradigms elicit a mismatch (MMN) response^4,5^, or sensory prediction error. According to the predictive coding framework and the model-adjustment hypothesis for MMN^6^, this error response is caused by a violation to the regularities, arising when sensory input does not match the prediction according to a learnt model^7^. The precision of this model varies according to the precision of context itself, such that a noisier environment will lead to the formation of more unreliable predictions than those formed in a stable environment^8^.

The MMN is robustly reduced in schizophrenia^9,10^, first episode psychosis^11^ and in individuals with high risk for schizophrenia^12,13^, hence suggesting that the ability to generate a precise model of the environment is reduced in schizophrenia and to some extent in the continuum of psychosis^14,15^. Recent predictive coding accounts put forward that the range of psychotic symptoms in schizophrenia can be explained by a failure to build precise beliefs or a model of the world^3^. The underlying processes of MMN generation involve both adaptation (neural habituation due to repeated standard sound stimuli) and prediction formation (guessing what might come next)^16,5^. It is unclear whether the consistent reductions of MMN in schizophrenia are caused by a failure of either adaptation or prediction, or both^17^. Adaptation and prediction formation are hard to disentangle with classical oddball paradigms because they typically evoke both processes simultaneously. Effective connectivity modelling attempts (i.e. inferring the dynamic effect one brain region has on another^18^), point to both adaptive and predictive processes being affected in schizophrenia^19^. In that study, dynamic causal modelling (DCM) was used to show that patients with schizophrenia have altered connectivity within the top-down connection from the right inferior frontal gyrus to the right superior temporal gyrus, as well as in the intrinsic connection (self-connection) within right primary auditory cortex. These two types of connections (intrinsic and top-down) are believed to reflect adaptation and prediction processes respectively, indicating that both processes were affected in that sample. We have further shown evidence that the same connections are altered in non-psychotic individuals with a genetic high risk for developing schizophrenia^20^. Such connectivity disruptions might explain the failure to build a precise (top-down) belief or model of the world. While the majority of previous studies on MMN in schizophrenia have been case-control studies assessing group differences, recent results indicate that a continuum perspective of psychosis is more powerful in explaining real-world functioning rather than a categorical approach as proposed in the Diagnostic and Statistical Manual of Mental Disorders (DSM)^21–23^. According to the continuum view, the major clinical symptoms reflect the degree of alterations in brain function resulting in functional abnormalities^21^. Hence, while the categorical approach is very useful for diagnostic purposes, the behavioural and brain alterations may not always follow this dichotomy and its aetiology might be best aligned on a continuum. An evident manifestation of this is precisely the MMN attenuation over the psychosis continuum^14^.

Here, we set out to investigate prediction errors with precision of the prediction manipulated to be either high or low in inpatients with a schizophrenia spectrum disorder (schizoaffective disorder or schizophrenia), inpatients with a psychiatric disorder but without psychosis, and healthy controls. We will adopt two approaches, a group comparison and a continuum approach along the dimension of psychotic-like experiences across all participants (N = 62, patients and controls) and parameterised by psychotic like experiences. The group approach will enable inferences on the specificity of connectivity changes in the schizophrenia spectrum group given our inpatient control group with a psychiatric diagnosis but without psychosis. The continuum approach will allow us to make inferences about how precision encoding and brain connectivity varies with the degree of psychotic-like symptoms irrespective of group membership (or diagnosis). We use a stochastic MMN paradigm previously validated in healthy controls^24^ that allows us to tap into predictive processes while mitigating adaptive processes, as well as to manipulating precision levels in the auditory environment. We hypothesise that the ability to encode the level of precision is reduced in the schizophrenia spectrum group compared to the two (non-psychosis and healthy) control groups. We expect this will be present both at the scalp level as well as in the effective connectivity. In addition, we hypothesise that the ability to encode precision decreases over the continuum of psychotic-like experiences across the whole sample.

## Methods

### Participants

62 participants took part in the study, with 20 inpatients with a schizophrenia spectrum disorder (SZS), 20 non-psychotic inpatients controls (NP), and 22 healthy controls (HC). Inpatients were recruited from the Monash Medical Centre, acute adult psychiatric inpatient facility. HC’s were recruited through the Psychology Research Participation Scheme (SONA), using Gumtree, and flyers distributed around the Monash Medical Centre. Participants provided written informed consent before taking part in the study. The inpatients were clinically assessed by their case clinician with respect to their capacity to consent to participate, before referral to the research study. Participants received monetary reimbursement for their time. This research was approved by the Monash Health Human Research Ethics Committee.

The positive and negative syndrome scale (PANSS)^25^ for schizophrenia was used to assess the symptoms of the two inpatient groups. In addition, the community assessment of psychic experience (CAPE)^26^ was administered to all participants in order to asses psychotic-like experiences across the continuum, including all three groups. The National Adult Reading Test (NART)^27^ was administered to get a proxy for IQ. Diagnoses for both groups of patients were made by the treating psychiatrist based on DSM-V criteria. Healthy controls were excluded if they had a history of a psychiatric or neurological disorder. Diagnoses within the NP group were: major depressive disorder; borderline personality disorder; posttraumatic stress disorder; or general/social anxiety. People in SZS group either had a diagnosis of schizoaffective disorder or schizophrenia. Groups were comparable with respect to sex ratio (HC: 12 males and 10 females, NP: 13 males and 7 females, SZS: 15 males and 5 females χ^2^ = 1.92, p = .38) and age distribution (HC: mean age = 34.09 years, standard deviation (SD) = 7.96 years; NP: mean age = 36.10 years, SD = 9.32 years; SZS: mean age = 34.30 years, SD = 7.33 years F_(2,61)_ = 0.37, p = 0.69). CAPE positive (CAPE+) scores differed overall between groups F_(2,61)_ = 3.049, p = 2.4*10^-5^). Post-hoc follow-up tests showed higher values for SZS than both NP (P = 0.020) and HC (P = 1.4*10^-5^) but there was no difference between HC and NP (P = 0.103). The SZS group had greater positive (t_38_ = −6.259, P = 3*10^-7^) and total (t_38_ = −4.691, P = 3.5*10^-5^) PANSS scores compared to the NP group. There was no difference on the negative (t_38_ = −1.422 p= 0.163) and general (t_38_ = - 1.588, p = 0.120) PANSS scores between NP and SZS. There was no difference in the approximation of IQ from the NART scores between the groups, see table 1. However, years of education varied significantly between the groups, which is counterintuitive to the IQ results. This may be due to variance between groups as to English being their first language. Chlorpromazine equivalent doses were calculated for the two inpatient groups to examine the effects of antipsychotic medications. A summary of the symptomatology and demographics can be seen in table 1 and in Supplementary Figure 1.

**Table 1:** Summary of demographic data. Illness duration had missing data for 5 NP and 3 SZS. WAIS IQ scores are predicted from the National adult Reading test. Abbreviations: WAIS = Wechsler Adult Intelligence Scale; CAPE = Community Assessment of Psychic experience; SD = standard deviation; Substance use disorders = SUD; Personality disorders = PD; Depressive disorders = DD; Anxiety disorders = AD; Schizoaffective = SZA; Schizophrenia = SZP; Somatic symptom disorder = SSD. Note the DD and SSD in the SZS group is a comorbidity with SZP.

|  | **HC (N=22)**<br>Mean (SD) | **NP (N=20)**<br>Mean (SD) | **SZS (N=20)**<br>Mean (SD) | **Statistic** |
| --- | --- | --- | --- | --- |
| **Sex** | Female: 10, Male: 12 | Female: 7, Male: 13 | Female: 5, Male: 15 | P = 0.380 |
| **Age (years)** | 34.09 (7.96) | 36.10 (9.32) | 34.30 (7.33) | P = 0.690 |
| **WAIS-R IQ estimate (from NART)** | 102.29 (7.60) | 103.47 (9.16) | 101.79 (7.21) | P = 0.800 |
| **Years Education** | 14.75 (2.31) | 12.75 (2.34) | 11.4 (2.11) | P = $5*10^{-5}$ |
| **CAPE positive** | 27.05 (6.44) | 32.15 (8.40) | 38.95 (8.02) | P = $2.4*10^{-5}$ |
| **PANSS positive** | N/A | 10.55 (3.85) | 21.20 (6.57) | P = $3*10^{-7}$ |
| **PANSS negative** | N/A | 11.40 (3.77) | 13.65 (5.99) | P = 0.163 |
| **PANSS general** | N/A | 30.85 (3.51) | 34.00 (8.14) | P = 0.121 |
| **PANSS total** | N/A | 52.80 (6.80) | 69.35 (13.70) | P = $3.5*10^{-5}$ |
| **Illness duration (years)** | N/A | 5.80 (6.08) | 9.58 (8.47) | P = 0.1627 |
| **CPZE** | N/A | 125.83 (220.87) | 583.70 (295.86) | P = $2.4*10^{-6}$ |
| **Diagnoses (including comorbidities)** | N/A | DD: 12<br>SUD: 8<br>PD: 6<br>AD: 4 | SZA: 7<br>SZP: 13<br>DD: 1<br>SSD: 1 |  |

### Stimuli

All participants engaged in a stochastic auditory paradigm in which participants listened to a stream of pure tones^24^. The tones were sampled from two different Gaussian distributions in log-frequency and centred at 500 Hz, see Fig 1A. The two distributions had either low variance (0.5 octaves above mean) or high variance (1.5 octave above mean). Tones were presented every 500 ms with a duration each of 50 ms and a rise and fall time of 10 ms. Within the random frequency stream, probe tones of either frequency equal to the mean (500 Hz) or 2 octaves above the mean (2000 Hz) were embedded pseudo randomly 10 % of the time. In what follows, we refer to the probe tone centred at the mean as the “standard” and the probe tone of 2000 Hz as the “deviant”. For an extensive description of the paradigm see Garrido et al (2013)^24^. Participants were asked to ignore the tones while performing a visual 1-back task, which entailed detecting repetitions of letters continuously presented, not coinciding with the sounds.

**Figure 1:**
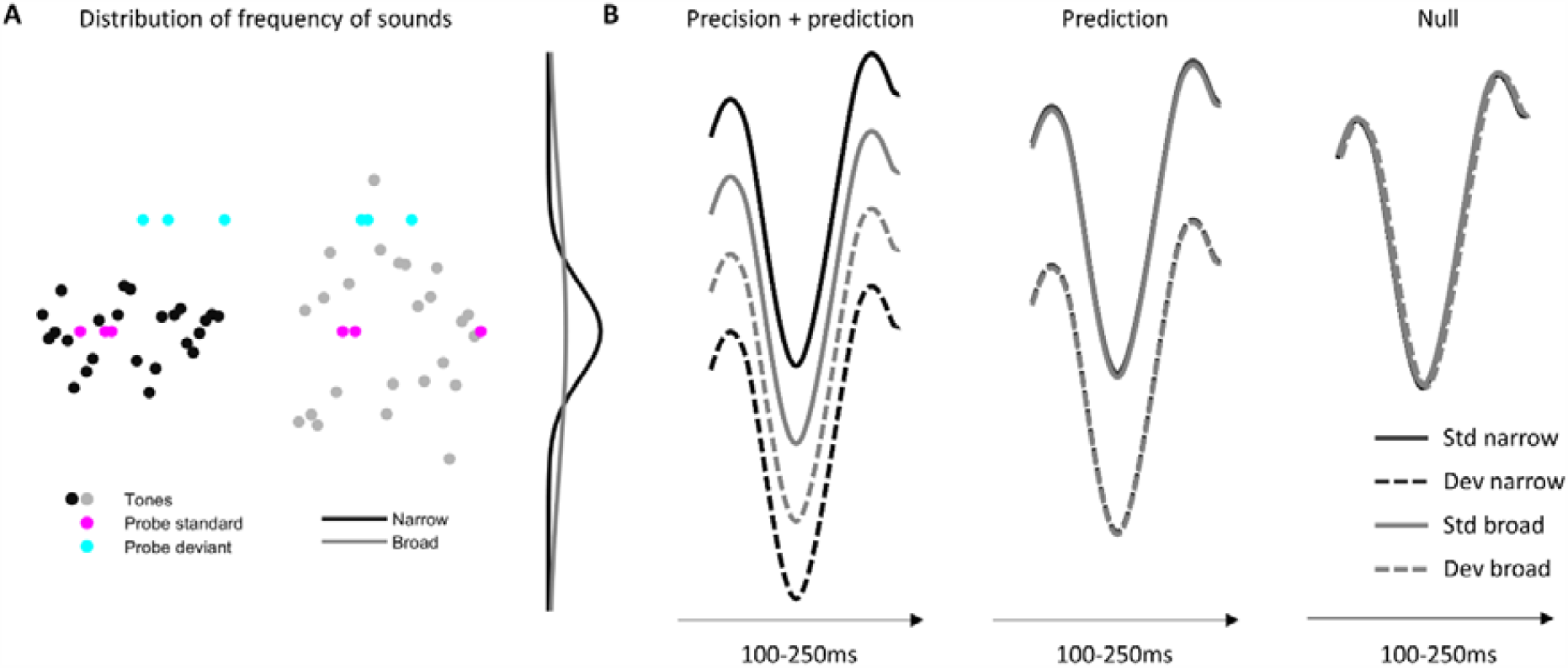
Auditory paradigm and the three models for ERP scalp data. A: The distribution of frequency of sounds are either following a narrow Gaussian distribution indicating high precision (black) or broad distribution indicating low precision (grey). Probe tones are indicated in turquoise for standards and magenta for oddballs. B: The precision + prediction model entails that both precision and prediction are encoded. The prediction model entails that a difference between standard and deviant tones are encoded but not the difference between broad and narrow context. Finally, the null model is representing that neither prediction nor precision is encoded.

### EEG recordings and preprocessing

EEG data was recorded using a 64 channel Biosemi active two with electrodes arranged according to the 10-20 system and a sampling frequency of 1024 Hz. All offline preprocessing was performed using SPM 12 (http://www.fil.ion.ucl.ac.uk/spm/), which included high and low pass filtering with a 5^th^ order Butterworth filter with a cut-off of 0.5 Hz and 40 Hz respectively. Data was epoched with a peristimulus interval of −100 ms to 400 ms, with baseline correction applied from −100ms to −5 ms. Finally artefact rejection was performed using a simple threshold technique rejecting trials if amplitudes exceed ±100 µV, and finally the signals were referenced to the average of all electrodes.

### Differences in MMN responses

Group differences in MMN responses were assessed using a one way repeated measures ANOVA testing for differences in mean values across groups with the factor precision (broad and narrow) as within-subject factor. Mean amplitudes of MMN calculated as the mean values of ±30 ms around the pooled group grand average peak were entered for each individual participant.

### Posterior probability maps – modelling the encoding of precision at the scalp level

We used Bayesian posterior probability maps on EEG data^28,29^ (at the scalp level) to test three different models: 1) prediction is formed, 2) both prediction and precision are encoded, 3) none are encoded - the null model. Bayesian maps enable the comparison of an arbitrary number of hypotheses (or explanations for observed neural responses), at each and every brain voxel (fMRI, or source level M/EEG) and/or in the scalp-time volume (scalp level), both within participants and at the group level. To compare the three models for event-related potentials (ERPs), we used Bayesian model comparison, on posterior probability maps^28,29^ as we have recently applied in Larsen et al., (2019)^16^. In this way, it is possible to infer across time and space which model is most likely. Epoched data were converted into scalp-map images of dimension 32×32 obtained using interpolation and smoothed using a Gaussian kernel specified by a FWHM of 8mm^2^ in the spatial dimension and 10 ms in the temporal dimension. Individual participant voxel-wise whole brain log-evidences were calculated using regressors describing the hypothesized relationship amongst the four conditions standard_narrow_, standard_broad_, deviant_broad_, deviant_narrow_ ([1 2 3 4] for the precision + prediction model, [1 1 2 2] for the prediction model and [1 1 1 1] for the null model), see Fig 1B. We did not test for the model only including precision and not prediction, given previous work demonstrating that prediction is indeed encoded in this particular paradigm^30,24^. Note that a model precluding prediction encoding would correspond to the absence of MMN. The log-evidence for each model were estimated using the variational Bayes first-level model specification^31^. Group level posterior probability maps were calculated using the random effects approach (RFX) for each model and each group. These probability maps can then be used to compare and select amongst the three different models for each voxel and time point.

### Dynamic causal modelling

Next, we used dynamic causal modelling (DCM) to investigate the underlying microcircuitry of precision encoding in the three groups. We were here interested in what specific connections encoding precision are modulated by the individual expression of CAPE+ both on a continuum and across the three groups. The network architecture underlying MMN has previously been established^32,20^ to include bilateral primary auditory cortex, the superior temporal gyrus and the inferior frontal gyrus. We made a fully connected network comprising intrinsic, forward and backward connections at all levels. We modelled the modulatory effect of precision with the prediction error response in the broad condition as baseline (low precision) and the narrow condition (high precision) as 1. All connections within the network were allowed to be modulated and this full model was inverted for each individual participant, see Figure 2.

**Figure 2:**
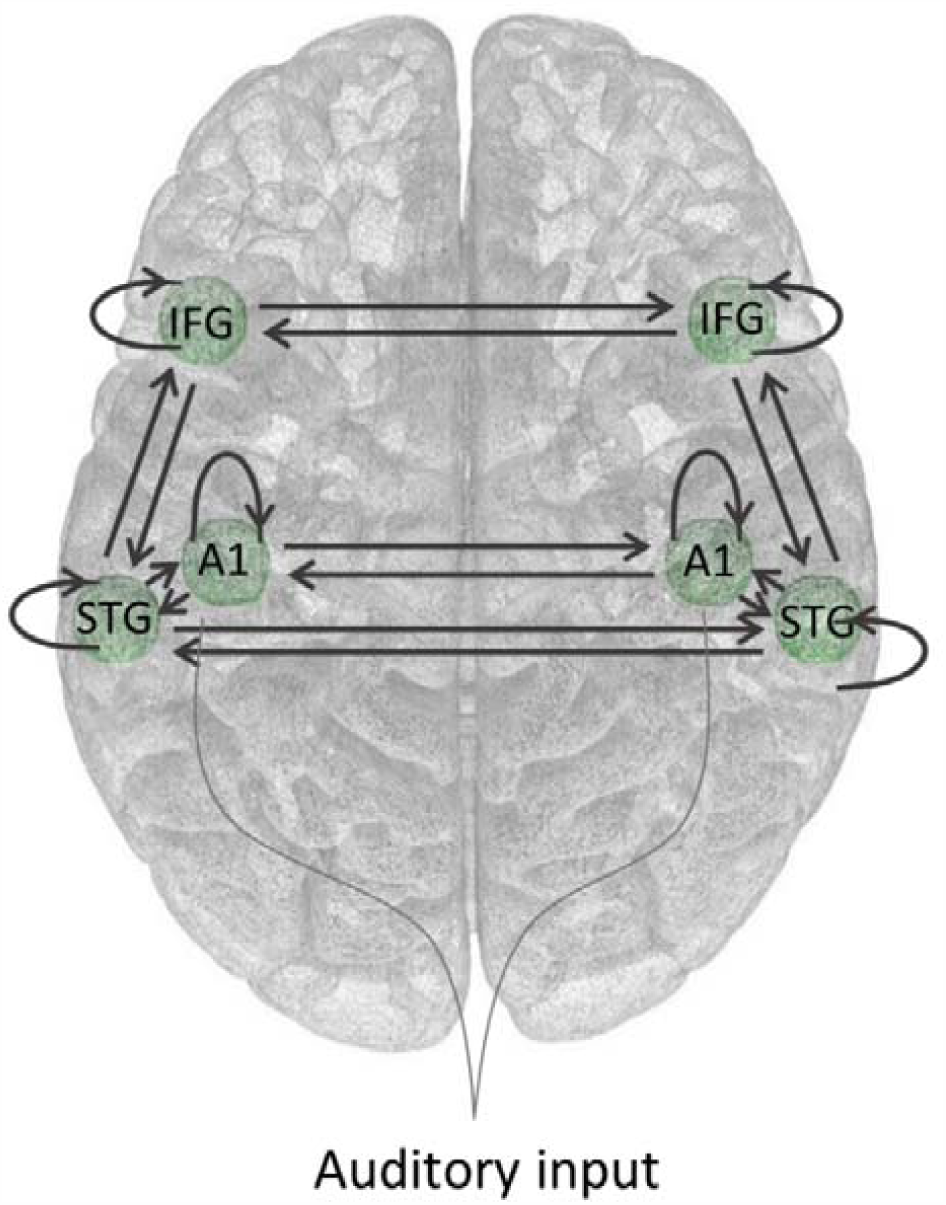
The full hierarchical DCM model, including bilateral inferior frontal gyrus (IFG), superior temporal gyrus (STG) and finally primary auditory cortex (A1). All connections are allowed to be modulated by the precision

### Modelling group differences in connectivity

To look into which connections modulated by precision of prediction error responses are modulated by group membership, we used a hierarchical model over the parameters as implemented in the parametric empirical Bayes framework^33^ in SPM12 (http://www.fil.ion.ucl.ac.uk/spm/). Here, we entered two regressors for the group membership, one modelling the main effect of being a patient (1 for the control group and −1 for both the NP and SZS group). The second regressor of interest modelled the difference between the two patient groups (0 for HC, 1 for the NP group and −1 for SZS group). Age and gender were entered as regressors of no interest and all regressors were mean centred. This design allowed us to tap into which connections were different overall between the controls and the two patient groups, and more specifically which connections were different between the two patient groups (i.e. specific to the schizophrenia spectrum). We report results that exceed 99% posterior probability, indicating very strong evidence that these connections are indeed modulated.

### Modelling changes in connectivity associated with CAPE+

To estimate systematic variations in model parameters on the continuum of CAPE+ scores, we created another second level model over the parameters. In this second level model, we entered CAPE+ scores as a regressor of interest together with age and gender as regressors of no interest. All regressors were mean centred. With this design we can estimate what connections in our DCM network change as a function of CAPE+ scores. We report with 99% probability on connections that had an association with CAPE+ (given the model tested).

## Results

There was no overall difference in the performance on the 1-back task when comparing d’ and reaction times across groups, see figure 2 in supplementary material. This rather easy task was intentionally chosen to keep participants engaged during the experiment, while avoiding potential confounds of task performance across groups.

### Precision is encoded in all groups

To test for differences in MMN responses between the broad and the narrow context, we took the mean around the peak for the MMN response for each group, Figure 3. MMN amplitudes differed between broad and narrow context across the three groups F_(1,59)_ = 25.188, p =5.1*10^-6^, showing a strong overall effect of precision. There was no difference in MMN responses between groups (F_(2,59)_ = 0.421, p = 0.658) as well as no interaction between group membership and the context (or precision) (F_(2,59)_ = 0.954, p = 0.391).

**Figure 3:**
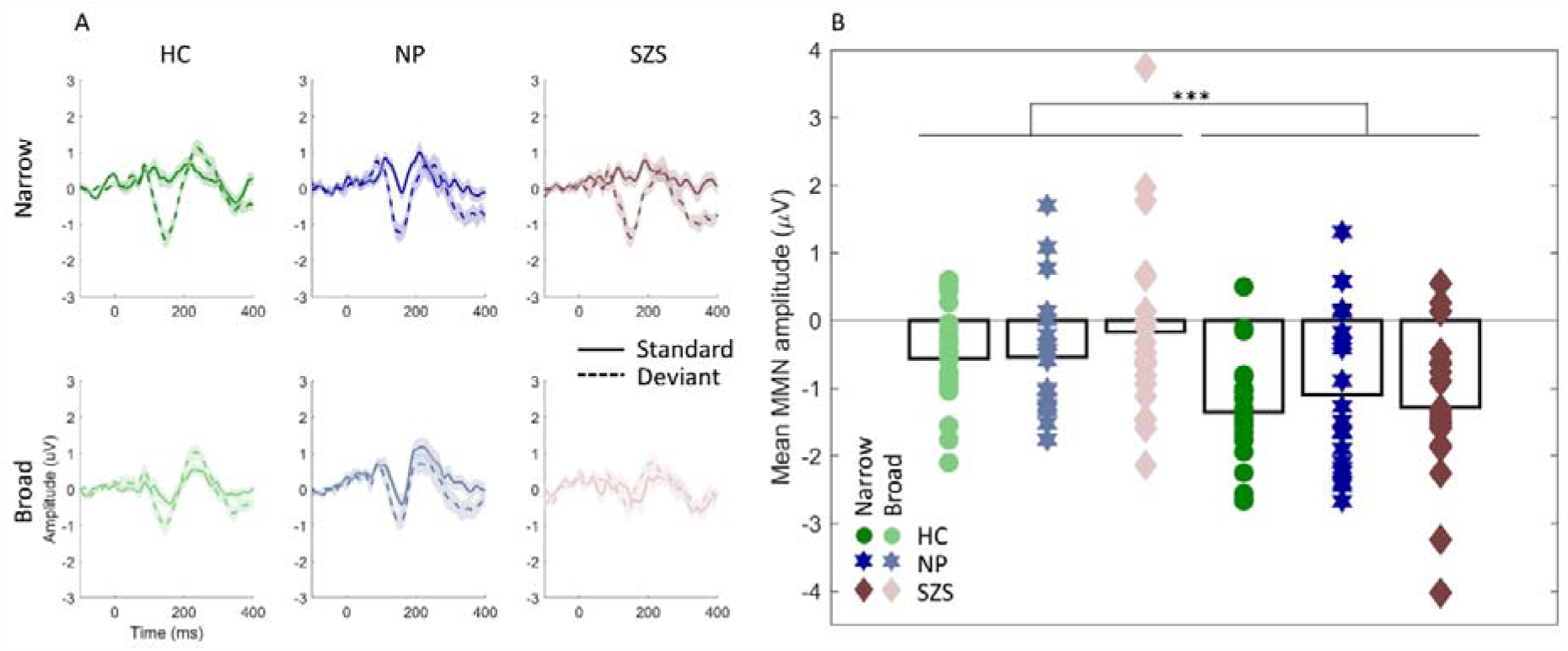
A: Responses to standard and deviant tones in the narrow (first row) and broad (lower row) context, NT in green, NP in blue and SZS in brown. Dark colours represents narrow context and light colours represents broad context. B: Mean MMN amplitude for the three groups in the two contexts, dots represents individual participants and bars group means, color-coding same as in A.

### Patients with schizophrenia spectrum disorder encode precision, but to lesser extent than non-psychotic inpatients and healthy controls

To answer the question of which of the three models were employed by each of the three groups, we plotted the exceedance probability as a function of time, summed across electrodes, Figure 4A. Overall, the null model expressing no difference between standards and deviants in the two contexts had highest exceedance probability in all time points except for the time window of the MMN response (100 - 200 ms), across the three groups. The model encoding precision (turquoise), as expected, showed highest probability in the time window of MMN, when prediction violations typically manifest. In that same window, HC and NP patients showed higher exceedance probability for the precision + prediction model than prediction only, whereas the SZS group showed equal probabilities across the two models. Hence, results indicate that during the time window of the MMN, HC’s and NP inpatients encode precision of prediction, whereas the SZS group show this to a lesser extent. To test whether this effect was significant between the groups, we entered the regressors of the precision + prediction model as a contrast into the GLM at the scalp level, allowing us to test for specific group effects in the precision encoding, Figure 4B. The SZS group showed reduced responses relatively to NT peaking at 248 ms in central-temporal channels, and at 104 ms in right fronto-central channels compared to the NP group. NP further showed an increased response compared to HC’s at fronto-central channels peaking at 107 ms, see figure 4B. No significant effects were found for the reversed comparisons.

**Figure 4:**
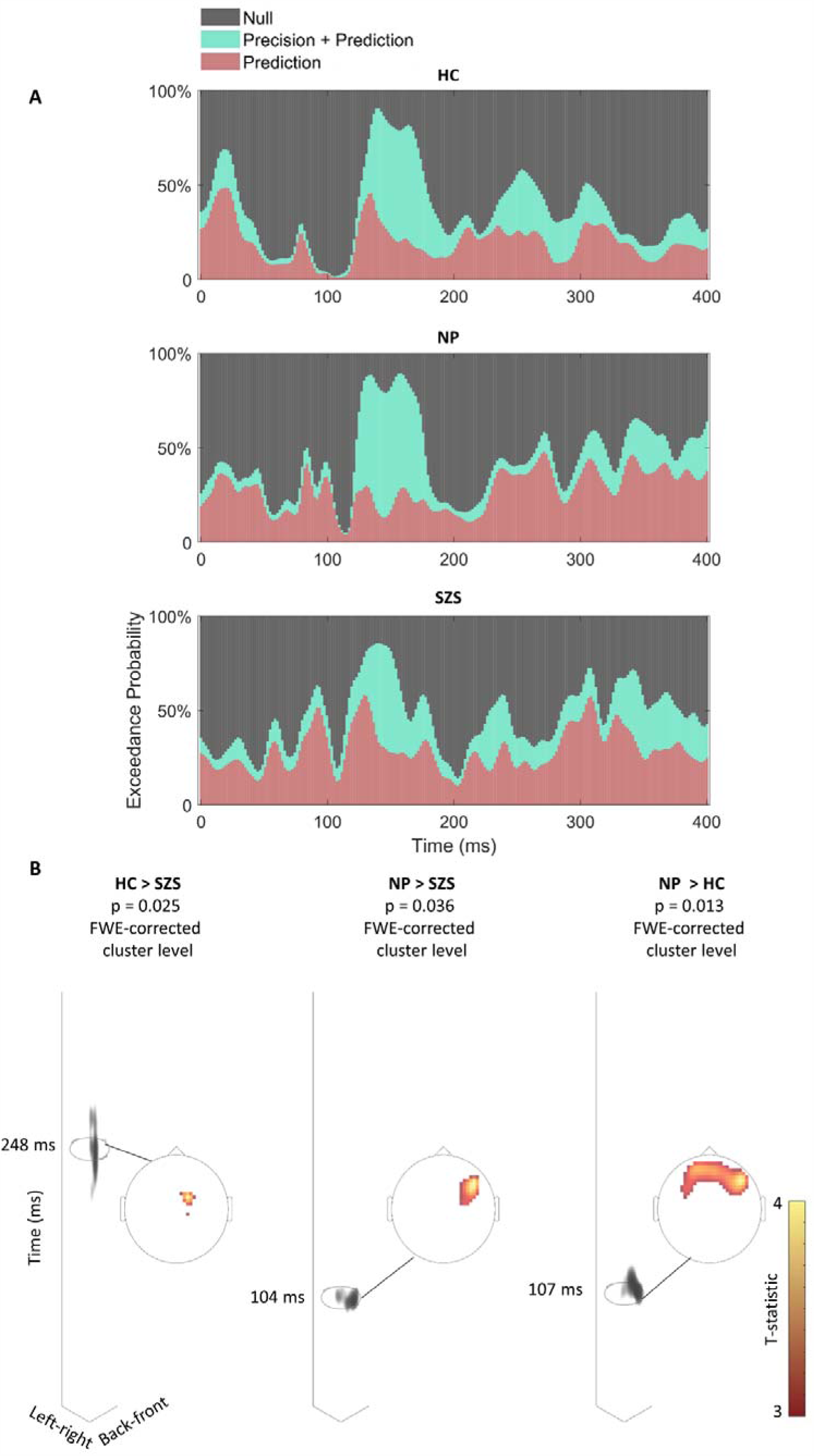
Precision and prediction are less encoded in schizophrenia spectrum disorder. A: Exceedance probability as a function of time, summed across space for the precision + prediction model (turquoise), prediction model (red) and null model (black). B: Regressors from the prediction+precision model were entered into the second level GLM to test for group differences for this effect across time and space. To the left statistical T-maps for HC > SZS is shown, middle is NP > SZS and finally to the right NP > HC. On the time axis it can be seen where in time there is significant activity for the given contrast as indicated with grey shading. The significant areas (over electrodes) for a given contrast are visualised within the topographic plot. The colour indicates a correspondence with the T-statistic above significance, with yellow being the strongest, see also Taylor et al., 2019^34^ for further description on this visualisation method.

### Brain connectivity is modulated by group membership

We asked which connections encoding precision differed between group membership. We found that the two patient groups overall had decreased lateral connectivity strength from left to right STG as well as top-down reductions from right IFG to STG when compared to the neurotypical control group. In comparison, the top-down connection from right STG to right A1 and the lateral connection from right to left STG was increased overall in the two patient groups compared to HC’s, see Figure 5. Critically, we found that five lateral connections differed between the two patient groups, hence indicating specificity to the schizophrenia spectrum. Specifically, connections from left to right IFG and right to left A1 were decreased in the SZS group compared to the NP group. Lateral connections from the right to left IFG, as well as bidirectional connection between left and right STG, showed increased strength in SZS compared to the NP group. We assessed any effects of medication on the connectivity parameters by means of Pearson correlations on all connections, which revealed greater connectivity from left to right STG for greater chlorpromazine equivalent scores (ρ = 0.422, p = 0.033). None of the other connections correlated with the medication dose.

**Figure 5:**
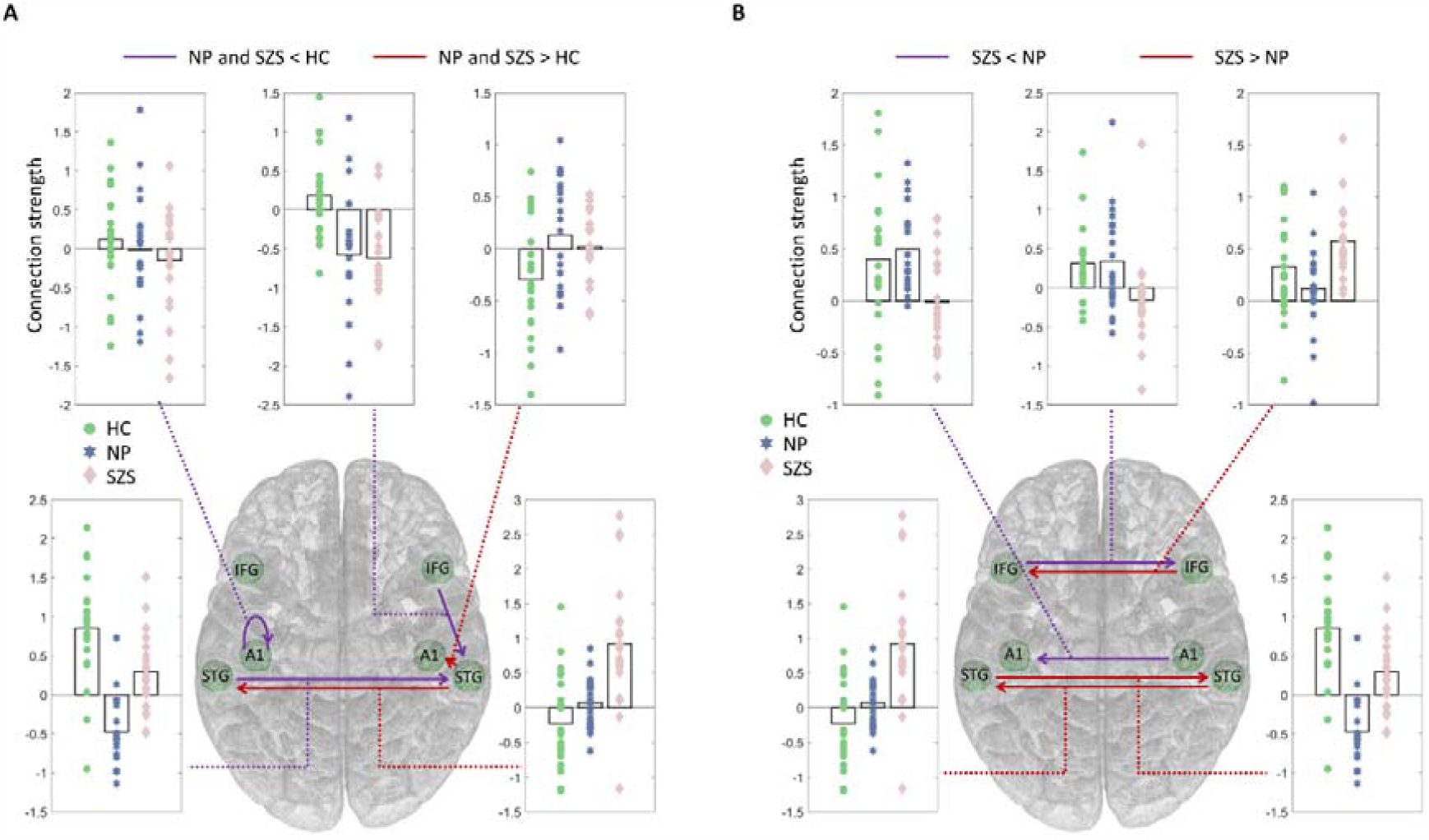
Connections modulated by group membership with an exceedance probability of at least 99% (very high evidence). A: main effect of patient, hence overall difference between HC and the two patient groups. Purple indicates connections where the two patient groups showed reduced connectivity when compared to HC, and red is where HC show less connectivity than the patient groups. B: connections showing differences between the two patient groups. Purple indicate connections where SZS show less connectivity than NP and red indicates SZS > NP. Note that the left to right STG connection is likely driven by medication effects. Dots correspond to individual participants in the HC group (green), NP (blue) and SZS (light red).

### Brain connectivity is modulated by psychotic-like experiences CAPE+

Most of the individuals in the SZS group, and in the NP group to some degree, had psychotic symptoms as measured by the PANSS. Across both patient groups there was a high correlation between CAPE+ and the PANSS positive symptoms score (ρ = 0.488, p = 0.001), indicating a high consistency between CAPE+ and PANSS positive symptoms. We therefore used the CAPE+ (instead of PANSS, which was only administered in the two patient groups) in order to align all participants on a continuum of psychotic-like experiences, including those in the healthy control group. By modelling the difference in contextual precision using the parametric empirical Bayesian framework in DCM, we were able to reveal which connections account for precision encoding and are modulated by the individual expression of CAPE+ across the whole sample. As mentioned previously, we report connections with associations above 99% exceedance probability. Lateral connections in both directions between IFG, as well as the top-down connection between right IFG and STG was negatively associated with CAPE+. The lateral connection from right to left STG as well as the bottom-up connection from right STG to right A1 were positively associated with CAPE+, see Figure 6. Hence, we see differences not only in the connectivity underlying precision encoding between the groups, but also individual connectivity alterations as a function of CAPE+ scores across the psychosis continuum. None of the connections associated with the CAPE+ scores were correlated with the chlorpromazine equivalent doses.

**Figure 6:**
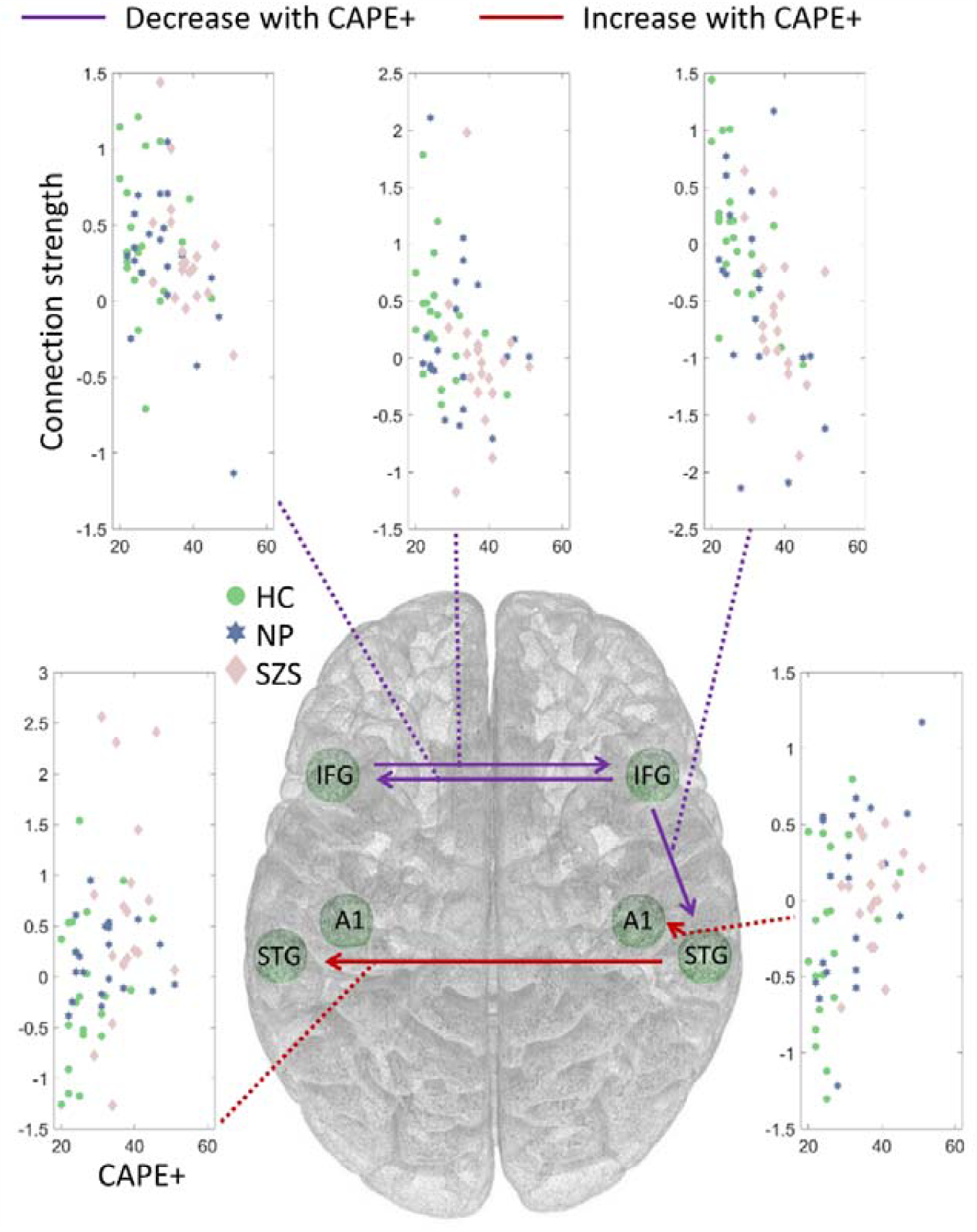
Connections modulated by individual differences in CAPE+ scores, showing decreases with CAPE+ (purple) and increases (red). Dots correspond to individual participants in the HC group (green), NP (blue) and SZS (light red).

## Discussion

Here we show that patients with schizophrenia spectrum disorder have alterations in the encoding of contextual precision that are associated with aberrant brain effective connectivity. Critically, we show that part of this connectivity alteration is specific to schizophrenia spectrum with the effect being present in the schizophrenia spectrum group but not the two control groups. We found not only group differences in the connectivity, but also connectivity changes across individuals on the continuum of CAPE+ scores. In addition, we showed that the schizophrenia spectrum group encodes both precision and prediction, albeit to a lesser extent than the respective non-psychotic patients and healthy controls. Medication effects were confined only to one connection linking left to right superior temporal gyrus.

We found group differences in the top-down connection from right IFG to STG. Further, this connection decreased with CAPE+. Perhaps not coincidently, this exact connection has previously been found to be weaker in schizophrenia^19^ and in non-psychotic individuals with increased genetic risk for schizophrenia ^20^. According to the predictive coding framework, top-down connections convey predictions, suggesting that as CAPE+ increases, distorted predictions are sent down the cortical hierarchy where precision is encoded. However, this connection did not show specificity to the schizophrenia spectrum group since no difference was found between this group and the NP group. In addition, we found that the two patient groups had decreased intrinsic connectivity within left A1, a connection thought to reflect adaptation^35^. This connection was found exclusively in the patient groups, hence not specific to the SZS group nor modulated by CAPE+. Therefore, considering our findings we suggest that this connection is affected more generally across psychiatric disorders or may be medication related, although there was no correlation with antipsychotic dose.

The SZS group showed reduced connectivity in lateral connections from left to right IFG and from right to left A1 as compared to the NP patient group suggesting that such reductions are specific to schizophrenia spectrum disorders. This is in line with previous results as it is robustly found that schizophrenia is associated with an interhemispheric disconnection^36–39^. Previous results on functional connectivity have shown that interhemispheric connectivity can distinguish schizophrenia patients from patients with depression, indicating specificity for schizophrenia^40^. Here we show evidence for specificity in the directed functional connectivity between hemispheres elicited by precision encoding in auditory stimuli. While we found reductions in connectivity that are specific to the schizophrenia spectrum, we also found some increases, namely in the lateral connections from the right to the left inferior frontal gyrus. An increase in the left to right superior temporal gyrus connections was found for the schizophrenia spectrum compared to the non-psychotic patient control group, although this (but not the former) appears to be driven by medication effects.

Somewhat unexpectedly, we found no evidence for MMN reductions in our SZS group. While reduced MMN in schizophrenia is very robust for duration oddballs, it is not the case for frequency deviants, as employed here. Indeed the effect size for MMN deficits in schizophrenia to frequency deviants are less than effect sizes for duration deviants^41^. In addition, the paradigm used in the present study is stochastic, meaning that it is not a sequence-based rule typically seen in auditory oddball paradigms. A recent meta-analysis suggested that effect sizes for MMN deficits in schizophrenia are reduced in auditory oddball paradigms with a more complex deviant type or pattern rule, as compared to standard oddball paradigms, where no complex rules are present ^41^. Both the use of frequency (instead of duration) oddballs and the complexity of the paradigm might explain the absence of the typically observed MMN reductions in the schizophrenia spectrum group. However, the connectivity results mentioned above, indeed show evidence for group differences, as well as differences across the continuum of CAPE+ scores, showcasing the usefulness of multivariate approaches (DCM) and Bayesian mapping as more sensitive methods for detecting group differences in brain activity than the traditional univariate EEG single channel analysis.

Here we demonstrate that while contextual precision is encoded in individuals with schizophrenia spectrum disorder it is so to a lesser extent compared to both non-psychotic patients and neurotypical control groups. However, we cannot conclude from this data if prediction is impaired, since the univariate analysis did not show reduced MMN for the schizophrenia group. We have previously shown that adaptive but not predictive processes are impaired in non-psychotic individuals with increased genetic risk for schizophrenia^16^. Future studies are needed to elucidate whether the reduction of MMN in schizophrenia is caused by impaired adaption or impaired prediction.

In conclusion, we show that connectivity underlying precision encoding is altered across the continuum of psychotic-like experiences. Critically, we show alterations (decreases and increases) in interhemispheric connectivity that are specific to the schizophrenia spectrum group and top-down fronto-temporal connectivity reductions that align on the continuum of psychotic-like experiences across both patients and healthy individuals.

## Supporting information

Supplementary matrial

## Acknowledgements

We would like to thank all included participants for their time in this study as well as the nurses in the psychiatric ward at the Monash Medical Centre, Adult Inpatient Psychiatric Facility.

## Funding

This work was funded by the Australian Research Council Centre of Excellence for Integrative Brain Function (ARC Centre Grant CE140100007), a University of Queensland Fellowship (2016000071), and a Women’s Academic Fund Maternity funding from Queensland Government, to MIG. OC is supported by the Australian Research council FFT40100807.

