## Supplementary matrial for "Aberrant connectivity in auditory precision encoding in schizophrenia spectrum disorder and across the continuum of psychotic-like experiences"

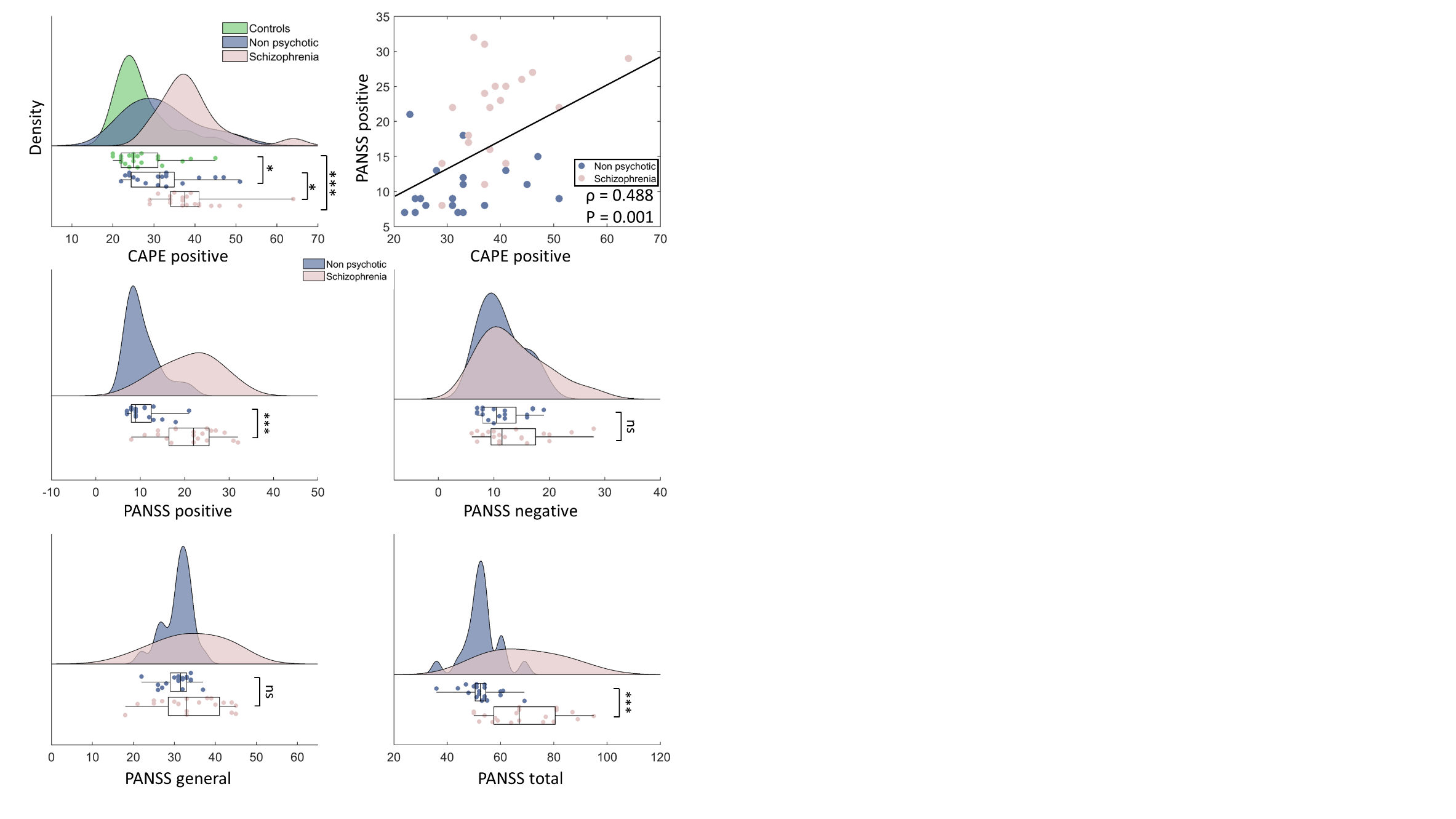

Figure 1: Distribution of the CAPE+ scores for the three groups, green is controls, blue non-psychotic and rose is schizophrenia. The four sub-dimensions for the PANNS (positive, negative, general and total) is shown for the non-psychotic (blue) and schizophrenia (rose). Top right shows correlation between self reported CAPE+ scores and PANSS positive scores for the schizophrenia and non-psychotic group.

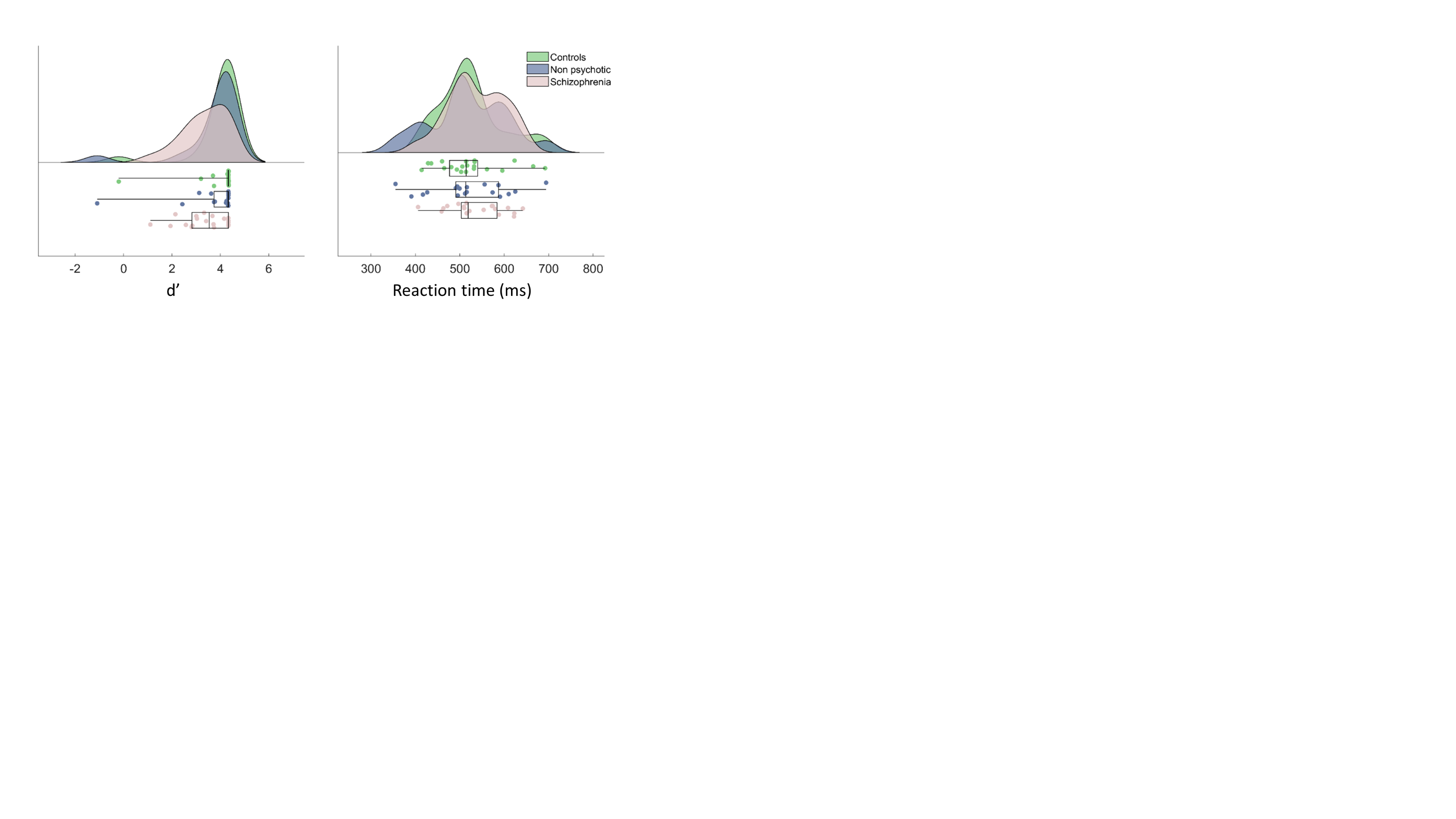

Figure 2: Results from the visual 1-back task, with d prime to the left and reaction times to the right. Again green is controls, blue non-psychotic and rose schizophrenia. There was no significant difference between groups on measures of d’ and reaction time.
